# A cryptography-based approach for movement decoding

**DOI:** 10.1101/080861

**Authors:** Eva L. Dyer, Mohammad Gheshlaghi Azar, Hugo L. Fernandes, Matthew G. Perich, Stephanie Naufel, Lee Miller, Konrad P. Körding

**Affiliations:** Dept. of Physical Medicine and Rehabilitation, Northwestern University, Chicago, IL; Sensory Motor Performance Program, Rehabilitation Institute of Chicago, Chicago, IL; Dept. of Biomedical Engineering, Northwestern University, Evanston, IL

## Abstract

Brain decoders use neural recordings to infer a user’s activity or intent. To train a decoder, we generally need infer the variables of interest (covariates) using simultaneously measured neural activity. However, there are many cases where this approach is not possible. Here we overcome this problem by introducing a fundamentally new approach for decoding called distribution alignment decoding (DAD). We use the statistics of movement, much like cryptographers use the statistics of language, to find a mapping between neural activity and motor variables. DAD learns a linear decoder which aligns the distribution of its output with the typical distribution of motor outputs by minimizing their KL-divergence. We apply our approach to a two datasets collected from the motor cortex of non-human primates (NHPs): a reaching task and an isometric force production task. We study the performance of DAD and find regimes where DAD provides comparable and in some cases, better performance than a typical supervised decoder. As DAD does not rely on the ability to record motor-related outputs, it promises to broaden the set of potential applications of brain decoding.

## Introduction

The aim of brain decoding methods is to infer the relationship between brain activity and a co-variate of interest. By measuring neural activity, one can infer what someone is viewing^1,2^, what word they are thinking about^3,4^, or their movements^5,6^. Decoding approaches can also be used to elucidate the link between external factors (stimuli) and the brain, which is essential for understanding how the brain encodes information. In all of these settings, training data are collected by simultaneously recording brain activity along with the covariates that we wish to predict.

In many situations, obtaining simultaneous recordings of both neural activity and the covariates of interest is challenging, expensive, or impossible^7^. This includes settings where complex task variables are hard to track and also cases where the user cannot generate training data (e.g., when a user is paralyzed). In these situations, there must be an alternative to applying a supervised approach to learn a mapping between neural activity and the underlying covariates that we wish to predict^8^.

To train a decoder without simultaneous measurements of neural activity and covariates of interest, we looked to cryptography for inspiration. When code breakers crack a cypher, they leverage prior information about the distribution of both individual characters and their joint statistics^9,10^. For example, the probability of observing a written ‘E’ is much higher than the probability of observing a ‘Z’. Using such information, Alan Turing and his Bletchley Park team cracked the World War II German Enigma code by exploiting the distribution of words known to appear in encrypted messages from their enemies. The key idea underlying this type of code-breaking strategy is to learn a mapping from the encrypted to decrypted text that produces a distribution with structure similar to what we expect based upon prior knowledge. Here, we ask if the same concept could be used to learn a mapping from neural activity to motor outputs.

We will highlight this concept with a movement example, where the aim is to decode the 2D velocity of a user’s hand from recorded neural activity. If the decoder is tuned correctly, the distribution of the decoded movement variables (Fig. 1a, orange) should roughly match the distribution of the kinematics of the actual movement^11^ (Fig. 1a, yellow). If the decoder is not correct, then its output would produce a different distribution (Fig. 1a, red). We will also do an analogous analysis for an isometric task, where the variable of interest is force, showing the applicability of our method to other motor-related variables. In cryptography, breaking a cypher is easier when the underlying signal is highly structured^9^. There is some evidence that natural movements, such as eating, running, and dancing, exhibit a great deal of structure and regularity^12^. The existence of such structure in natural movements suggests that deviations from the typical movement distribution could be statistically detected. Aligning the distribution of predicted movements with a prior over movements may provide a way of relating neural activity and movement without the need to measure them simultaneously.

**Figure 1:**
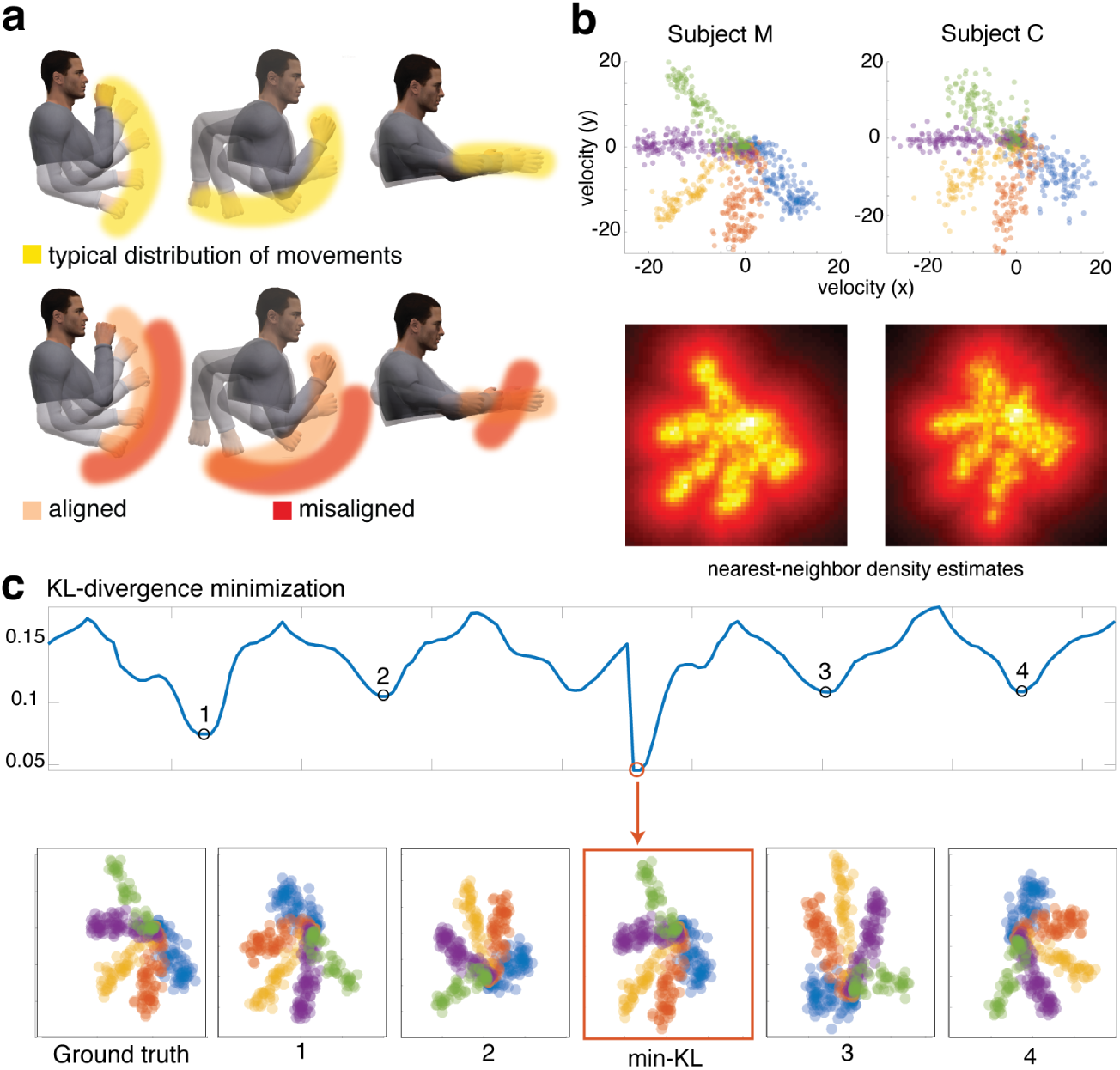
Decoding structured movements with distribution alignment. In (a), we show a schematic of hand trajectories of different tasks. For each task (feeding, running, reaching), the hand moves along a subspace within the space of all 3D movements. The distribution of hand movements for each task is highly structured and anisotropic and varies across different tasks. In (b), kinematics data from Subject M (left) and Subject C (right), displayed as scatterplot data with the target directions displayed as different colors and on the bottom, the corresponding heatmaps (distribution estimates). In (c), we demonstrate the idea behind distribution alignment on two permuted sets of training and test kinematics from Subject M. The KL-divergence is displayed as a function of the rotation angle used for alignment in 2D. The correct solution (ground truth), min-KL predicted solution (R^2^ = 0.996), four non-optimal rotations/reflections are displayed. A prior movement distribution from the same subject is shown in (b). Note that due to the fact that this reflections of this task are roughly equivalent, the first local minima (1) of the KL-divergence is a flipped-version of the correct solution.

Here we introduce an approach for movement decoding called distribution alignment decoding (DAD), that exploits the structure of natural movements to find a mapping between neural activity and movement. Our approach leverages the fact that low-dimensional representations of populations of motor neurons have the same shape (distribution) as the motor outputs ^13,14^. Thus, in order to find a decoder with output that matches a prior distribution of movements, we only need to find a low-dimensional representation of the neural data that matches this movement distribution. We do this by learning a decoder that minimizes the deviation (KL-divergence) between the distribution of projected (low dimensional) neural activities and kinematics.

We applied DAD to neural datasets collected from the primary motor cortex (M1) of three non-human primates (NHPs) performing either a reaching task or a task to produce isometric force about the wrist. When the covariates of interest are linearly embedded in the neural activity, DAD can be used to predict motor outputs without knowing the correspondence between training and test data. Our results suggest that it is possible to decode brain activity without the need to observe the covariates that we want to predict. Thus our approach can also be used to decode the neural activity of one subject based upon distribution of motor outputs collected from another subject— this is useful in settings where we have no previous motor data from a subject. Lastly, when a small amount of supervised data is available, we can combine the outputs of DAD and the supervised decoder to improve supervised approaches.

## Results

DAD projects the neural data into a low-dimensional space, and then finds a mapping of the resulting data into movement space so that the resulting distributions are as similar as possible. To evaluate its performance, we measured populations of neurons in the arm and hand areas of the primary motor cortex (M1) from three NHPs while they performed different motor tasks. We studied data from two motor tasks: a standard reaching task (Subjects M and C) and an isometric wrist force production task (Subject J). In addition, we collected samples of the covariates of interest in both tasks. In the case of the reaching task, we recorded the position and velocity of each subject’s hand. For the isometric task, we measured the forces about the wrist. To produce non-isotropic datasets with meaningful structure, we subsampled our data to remove a few target directions (see Fig. 1 for details on this choice). This way we have a non-isotropic movement distribution that we can align with the equally non-isotropic distribution of neural activities to test the performance of DAD.

The kinematics of the reaching task, as performed by different subjects (M and C), showed high degrees of similarity when aggregated over a large number of trials (Fig. 1b). We computed the distribution of both subject’s movements using a non-parametric nearest neighbor approach for density estimation and found that the distributions are extremely similar. The similarity between the distribution of different subjects’ movements suggests that our approach could be applied to decode neural data using training kinematics from another subject, which can be useful when no supervised training data is available for a particular subject.

To confirm that our method can, in principle, align low-dimensional data, we tested our approach on the 2D kinematics data from subject M (Fig. 1c). Our alignment procedure computes the KL-divergence between the training distribution and a transformed version of the target distribution. DAD returns the alignment that minimizes the KL-divergence between the training and test. The resulting alignments provide decoders with good prediction accuracy (as measured by R^2^). Thus the KL-divergence provides a measure for aligning two distributions and can be used to solve our alignment problem.

Distribution alignment is relatively easy when the training and test distributions have similar structure. However, low-dimensional embeddings of neural activity can be quite noisy, therefore it is important to first verify that the task structure is apparent in the embeddings. We applied a variety of linear and nonlinear dimensionality reduction techniques^15,16^ to embed the neural data (see Methods). In the case of both subjects M and J, we found certain directions of movement (M) and force production (J), that produce neural embeddings that lie on a cone in 3D (Fig. 2c). Moreover, we find that the structure of the individual movement directions is also preserved in these embeddings. When we applied the same approach to neural data collected from Subject C, we couldn’t find any discernible structure that could be used to align it with the target distribution. As we highlight in the discussion, the failure mode of DAD is typically the failure of dimensionality reduction algorithms to find the dimensions that contain the relevant signals. We therefore did not use Subject C’s neural activity for decoding, however, but we did use this subject’s kinematic data as a prior over movement. Our analysis suggests that dimensionality reduction can be used to first embed the neural data into a space where it can then be more easily aligned.

**Figure 2:**
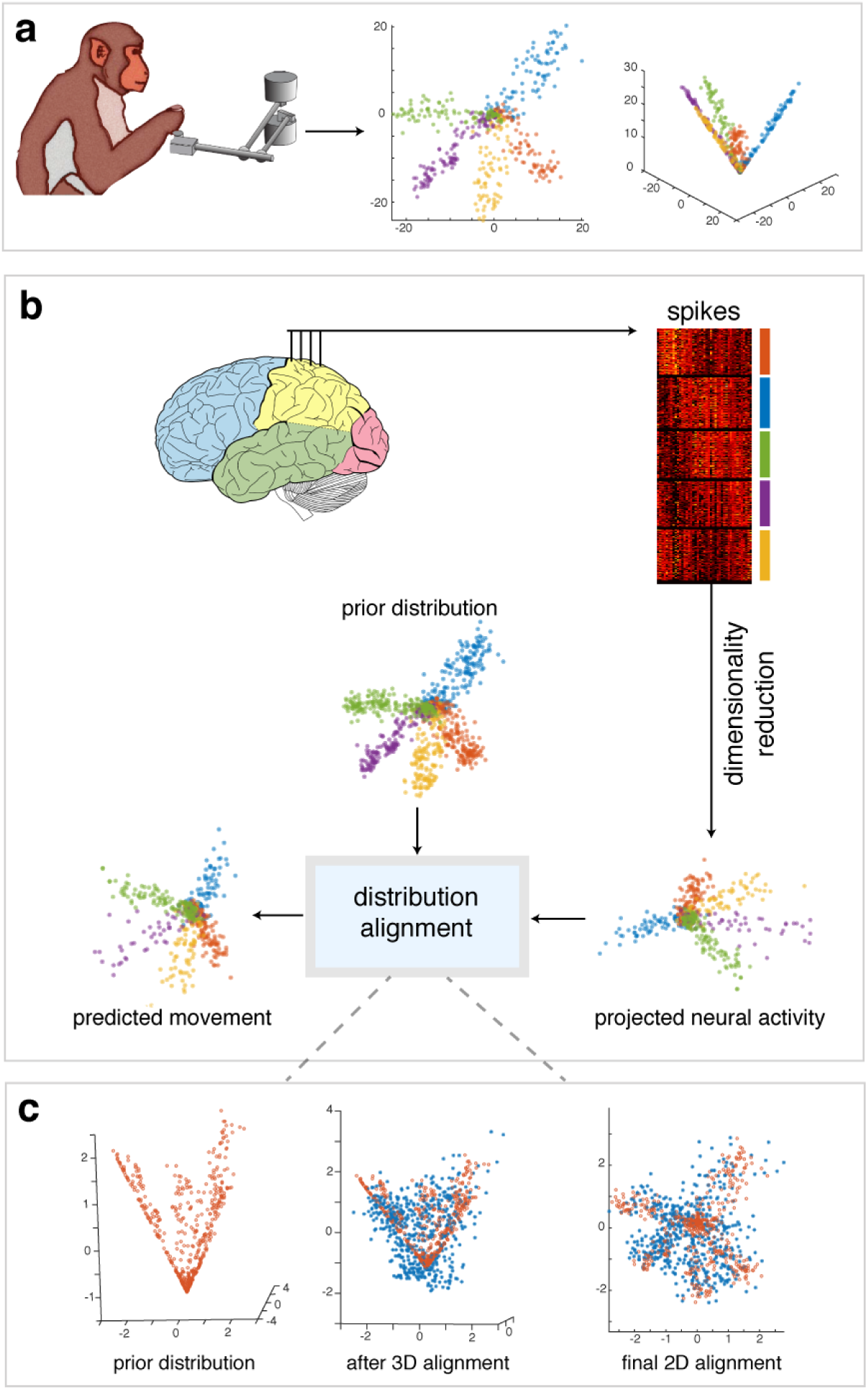
Overview of Distribution Alignment Decoding (DAD) Approach. In (a), we depict a subject carrying out a reaching task and show the resulting 3D kinematics associated with this task. The subject’s movements produce datasets with the (top) 2D kinematics of the task and (bottom) 3D movement data when augmented with speed as a covariate. (c) Neural activity in the form of spikes are mapped into a 2D predicted-movement space using factor analysis. DAD then finds an affine mapping that aligns the projected neural activity (embedding) with the prior distribution of kinematics. (d) We show the results of our alignment procedure in 3D, (left) the target distribution (red *◦*), (middle), the aligned neural data (blue ∗) overlaid on the target (mid), and the aligned distributions when projected into 2D.

### Performance on reaching tasks

To start, we analyzed neural data from Subject M during a reaching task. We explored three different training scenarios to test DAD: (i) a *within-subject* setting, where all of the training kinematics are from Subject M (DAD-M), (ii) a *combined-subject* setting, where the training kinematics contains samples from both M and C (DAD-MC), and (iii) an *across-subject* setting, where the training data is only from C (DAD-C). As DAD is relatively insensitive to the specific training distribution, we find that the alignments we obtain for the across- and combined-subject setting are often similar to the within-subject case (Fig. 3a). However, when there is little training data, we find that adding training data from different subjects can improve the performance of the decoder (Fig. 3b).

**Figure 3:**
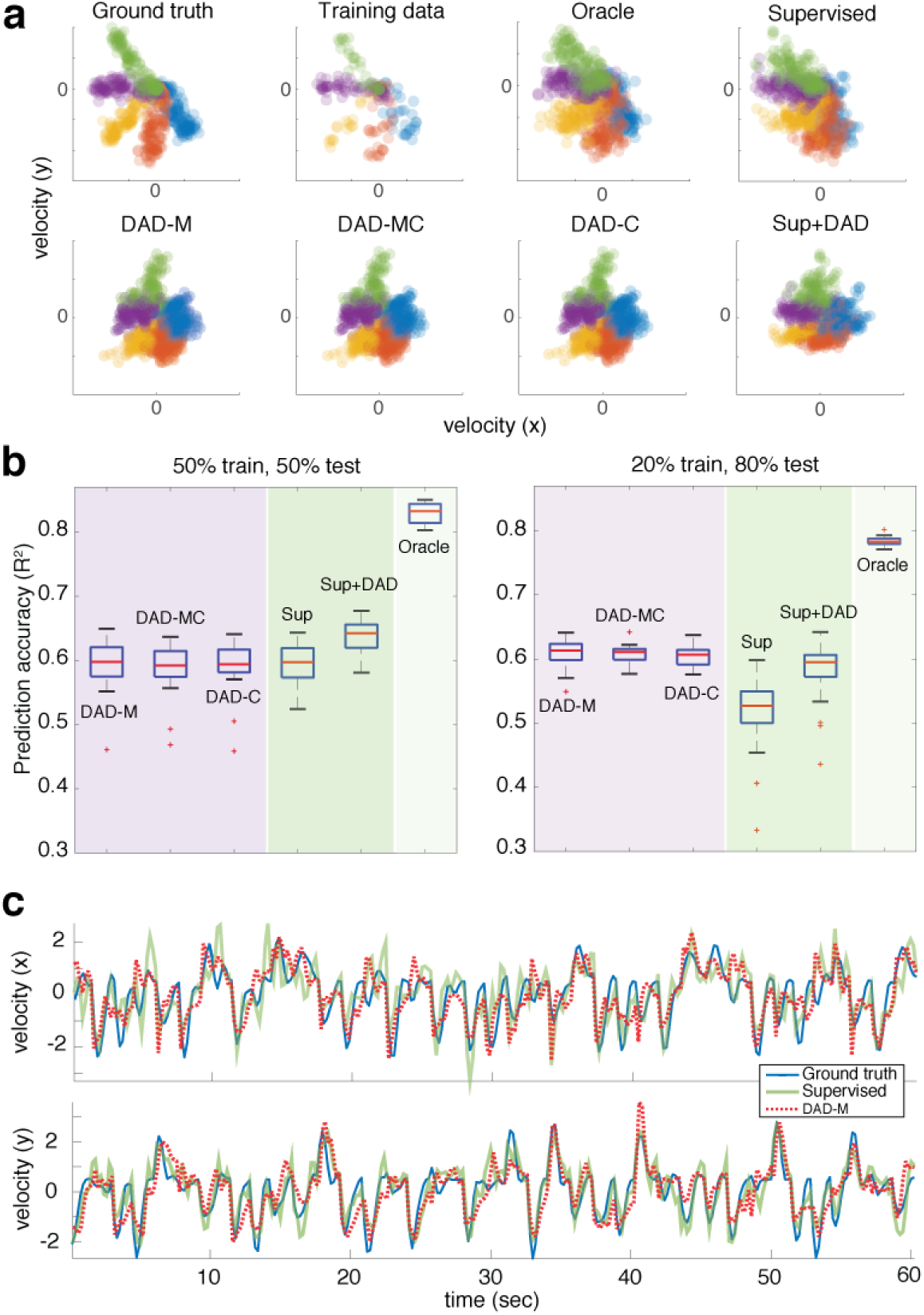
Comparison of decoding performance for reaching tasks. We compare the performance of DAD with a supervised decoder (Sup) trained on simultaneously collected neural and kinematics data and an oracle estimate (optimal linear 2D decoder). In (a), we show the result of DAD-M (train and test on M), DAD-MC (train on M,C and test on M), and DAD-C (train on C and test on M), when 20% of the dataset is provided for training. The prediction accuracy for DAD in all three cases is R^2^ = 0.627. The supervised decoder obtains a R^2^ = 0.572 and the oracle estimate is R^2^ = 0.781. In (b), we provide performance comparisons for supervised approaches (in shaded green regions) and DAD (in shaded purple regions) for different amounts of training and test data. We display boxplots of the R^2^ values for DAD, a supervised decoder (Sup), the average between the predictions of Sup and DAD-M (Sup+DAD), and the oracle, as we increase the amount of test data. Each boxplot shows the results over 20 trials, the median is displayed as a red line in each box, the edges of the box represent the 25% and 75% percentile, the whiskers indicate the range of the data not considered an outlier, and the outliers are displayed in red (x). In (c), we show the time-varying decoding performance of DAD-M and the supervised decoder (Sup) overlaid on the ground truth kinematics (200 ms time bins).

To compare DAD with a supervised decoder (Fig. 3b-c), we randomly partitioned the data into training and test sets and give both decoders access to both. Before seeing the test set, DAD is only provided the kinematics from the training set, while the supervised decoder has access to both movement kinematics and neural activity. During the test phase, DAD uses the training kinematics and the neural test set to learn a decoder. For our comparisons, we trained a supervised decoder with 10-fold cross-validation to fit the regularization parameter (see Eqn. 2 in Methods). In addition to a supervised decoder, we also compare with an oracle that finds the best possible 2D linear embedding that can be obtained for the test set. It is important to note that the so-called oracle decoder is impossible to obtain in any practical situation and only serves as a measure of the upper bound over the set of possible linear decoders.

We studied the performance of DAD as we varied the relative proportion of training and test samples. In our evaluations, we applied DAD to neural data from Subject M, using kinematics training datasets from the same subject (M), another subject (C), and from both (MC). We observe an interesting asymmetry in the performance of DAD versus supervised methods due to the way they are trained. In particular, we find that when the decoder is given access to a small amount of training data, DAD consistently outperforms supervised approaches and has smaller variance in its prediction accuracy (Fig. 3b). This is due to the fact that when there is little training data, DAD can still use the test data to improve its final alignment. However, as we increase the amount of training data, supervised approaches perform on average, slightly better than DAD. We average the outputs of DAD and the supervised decoder (Sup+DAD) and find that the combination often improves over the individual performance of each. These results are further confirmed by examining the time-varying decoding of the kinematics (over 200 ms time bins) in both the x and y directions (Fig. 3c). Our results suggest that DAD can be used in settings where little training data is available and in across-subject settings, where no previous movement data is recorded from a subject.

### Performance on isometric force production tasks

To test the performance of DAD on a different motor task, we applied it to neural recordings from another subject (Subject J) performing an isometric force production task. The subject’s hand was fixed and thus the measured covariates that we aim to decode are the 2D forces applied at the wrist. When we isolated the neural activity to time points right after the ‘Go’ cue (indicating that the subject should begin force production), we observed clear embeddings of neural data (Fig. 4a) that resemble the task structure. However, the size of the dataset is much smaller, and thus our embeddings are not as stable across multiple random partitions of the dataset. Our results for Subject J suggest that certain directions of force-production in an isometric task can also be decoded using DAD. DAD thus holds promise for a variety of motor tasks.

**Figure 4:**
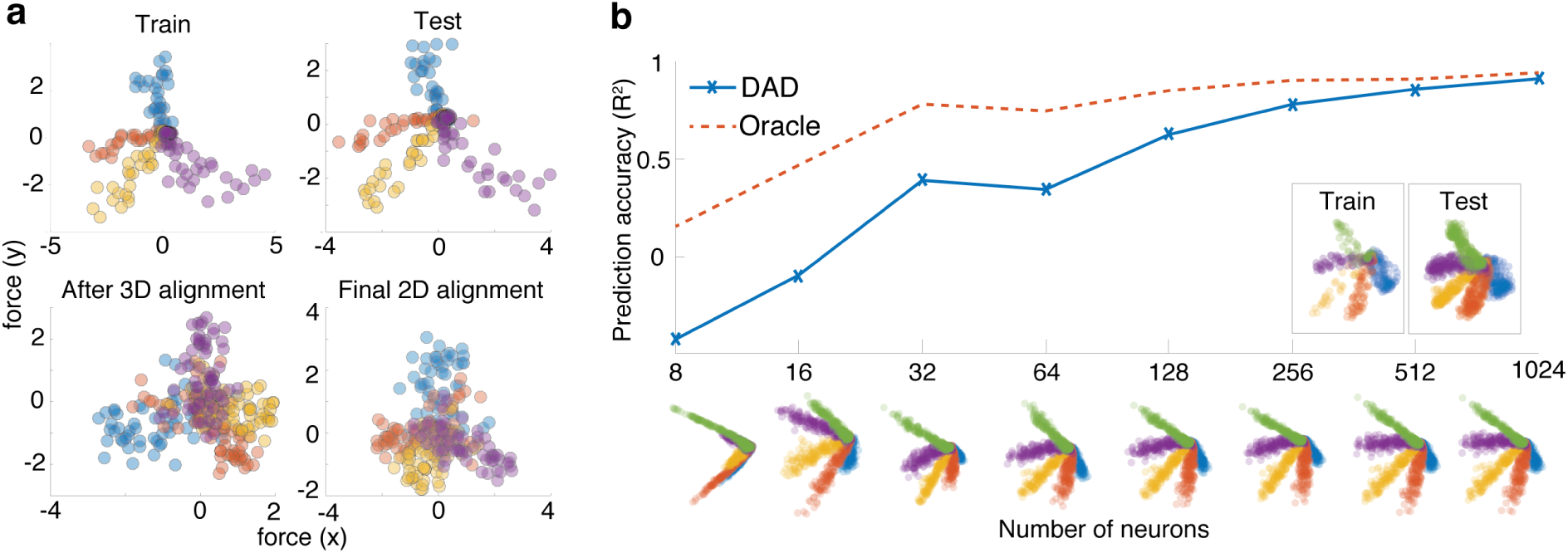
Prediction accuracy increases with population size. In (a), we show the results of DAD on an isometric task, where we record forces applied to the wrist to move a cursor to different targets. Along the top row, we display the training kinematics (used to estimate the prior distribution) and the test kinematics (ground truth). Along the bottom row, we show the output of our cone alignment procedure (3D embedding) and our final 2D estimate (R^2^ = 0.51). We observe that DAD indeed finds the correct 3D alignment and then the final 2D rotation to align the datasets. In (b), we visualize the accuracy of DAD on synthetic data as we increase the numbers of neurons in our model. We display the trimmed mean of the R^2^ values (the top and bottom 10% of trials are removed) obtained for DAD and the oracle, averaged over 100 trials. Below, we show examples of alignments obtained as we increase the number of neurons in our synthetic model. Examples of training and test kinematics are also shown.

### Synthetic experiments

In some of our experiments with real data, the low-dimensional projections of neural data become (nonlinearly) warped (Fig. 3a). Thus, to explore the relationship between the embeddings and the size of the neural population, we create data generated from a population of synthetic neurons with bio-realistic tuning properties. More concretely, we create synthetic data by generating a population of neurons with tuning curves drawn from a distribution of parameters designed to match the real datasets (see Methods).

Using this synthetic model, we studied the impact of both the number of neurons and time points used for dimensionality reduction and alignment (Fig. 4b). We compare the performance of DAD with that of the oracle decoder which has access to both the neural activity and kinematics of the test set. As expected, as the number of neurons increases, the performance accuracy increases as well. For this model, DAD performs as well as the oracle (best 2D linear fit) for populations with roughly 1,000 neurons. This model appears to align nicely with our results on real data; for populations of 100-200 neurons (typical recording sizes), the average R^2^ is around 0.6-0.7 (R^2^=0.62 and R^2^=0.78 for d = 128 and 256 neurons, respectively). Thus this suggests that with recordings from even larger populations, distribution alignment will become even more accurate.

## Discussion

In this paper, we introduced DAD, a cryptography inspired approach for brain decoding. In the context of motor decoding, we show how DAD can be used to learn a decoder by minimizing the divergence between the distribution of motor outputs and a projected distribution of neural activity. Thus, instead of requiring access to simultaneous recordings of neural activity and motor outputs, our approach relies on the structure and regularities of movement and the fact that this structure is preserved in low-dimensional projections of neural activity of the motor cortex. We find that the algorithm works well on simulated data, as well as two datasets collected from motor cortex.

One assumption that is integral to our approach is that the variables of interest, e.g. the low-dimensional velocities or forces, appear in the set of the first few components of a dimensionality reduction algorithm. We initially assumed this would be true, as cosine tuning is one of the most frequently described properties of motor cortex activity ^17^. However, this assumption is not true for many datasets, and hence DAD can not be applied. If motor cortex primarily deals with movement vectors, why do they not consistently show up in the first principal components? Despite actively analyzing the data, we can not currently answer this question. This is a very interesting finding as it suggests that there are important features of neural activity that we do not understand. A better understanding of the relevant variables could make DAD work dramatically better and for more datasets.

Our approach assumes that there exists a linear transformation between projected neural activity and the distribution of movements. However, this assumption often does not hold in practice ^18^. This is likely why we observe that the fine-scale structure of the predicted kinematics within each target direction is warped (Fig. 3a). An exciting line of future research would be to extend our approach to incorporate non-linearities. One possibility would be to explore the use of mixture models^19^ to model the low-dimensional structure of the data, instead of using a single subspace model with PCA or factor analysis. Kernel methods^20^ could also be used to lift the neural data to a higher-dimensional space where there exists a linear relationship between the neural activity and the low-dimensional kinematics. With approaches that address the non-linearity of the data, the underlying structure of movement contained in the neural data can be preserved, thus leading to an improvement in the prediction accuracy of the decoder.

To guarantee a unique solution for the problem of distribution alignment, the distribution of movements must be anisotropic (no rotational or reflection symmetry). However, in many laboratory tasks such as center-out-reaching tasks, this assumption is often violated. Here, we circumvent this problem by subsampling the dataset. This introduces asymmetries into the motor task and, in turn, guarantees the uniqueness of the resulting solution. While requiring non-isotropic movements is clearly a drawback of our approach, natural movement tasks^11,21^ (Fig. 1a) are more likely to produce non-isotropic distributions. Moreover, in the case of isotropic distributions where multiple alignments exist, we can potentially use prior knowledge of the decoder or feedback from the user, to rule out incorrect alignments. Since natural movements typically exhibit asymmetries, we expect that our approach can be applied to decode motor variables in a wide range of tasks.

While our results demonstrate that DAD can achieve comparable performance to that of a regularized linear decoder trained with cross-validation, state-of-the-art decoders use additional information to improve the decoder’s performance, e.g. modeling non-linearity in neural firing rates (e.g., Poisson models)^18,22^, target information^6^, smooth temporal structure^23^, and the drift of neural properties^24,25^. Since we do not leverage such improvements, using additional structure is likely to improve the performance of a supervised approach. Similarly, the performance of DAD could be significantly improved using the same ideas. One such improvement would be to incorporate temporal structure by stacking recordings from consecutive time steps together and then aligning the distributions of these augmented vectors. We expect that by incorporating temporal structure, as well as other insights gleaned from studies of the low-dimensional structure of neural populations^26–28^, the performance of DAD will be further improved.

We solve the decoding problem by exploiting the known structure of movements to learn a decoder that produces outputs that are aligned with this structure and thus demonstrate one way in which cryptography might applied to brain decoding. The approach we used here is quite simple, but we imagine that more sophisticated code breaking strategies could be used in this and other brain decoding scenarios (beyond movement). Thus, ideas from cryptography and distribution alignment promise to enable a broad range of approaches into brain decoding and new insights into how to crack the neural code.

## Methods

### Data collection

Neural and behavioral data were collected from three rhesus macaque monkeys (we refer to them as Subject M, C, and J). All surgical and experimental procedures were approved by the Northwestern University Animal Care and Use Committee, and were consistent with the National Institutes of Health Guide for the Care and Use of Laboratory Animals.

In the first experiment (Subject M and C), the subjects performed a standard delayed center-out movement paradigm (reaching experiment). The subjects were seated in a primate chair and grasped a handle of a custom 2-D planar manipulandum that controlled a computer cursor on a screen. The subject began each trial by moving to a 2 × 2 × 2 cm target in the center of the workspace. The subject was then required to hold for 500 – 1500 ms before another 2 cm target was randomly displayed in one of eight outer positions regularly spaced at a radial distance of 8 cm. For Subject M, this is followed by another variable delay period of 500 – 1500 ms to plan the movement before an auditory ‘Go’ cue. The sessions with Subject C omitted this instructed delay period and the ‘Go’ cue was provided when the outer target appeared. The subjects were required to reach to the target within 1000 – 1300 ms and hold within it for 500 ms to receive an auditory success tone and a liquid reward.

In the second experiment (Subject J), the subject performed a center-out isometric task. The hand was placed in a box attached to a 6-axis force-torque sensor (JR3, Inc, Woodland, CA), and the subject applied force about the wrist to move a cursor on a screen. The position of the cursor was linearly related to the applied force. The subject began each trial by keeping the hand relaxed so that the cursor rested on a start target on the screen. After staying in that target for 200 – 1000 ms, a second target appeared and the subject was required to apply force to reach this target within a period of 2000 ms. There was no delay period in this task, therefore the subject could go to the target as soon as it appeared. The subject was then required to hold within this target for 500 ms to receive an auditory success tone and liquid reward.

After the subjects received extensive training in each task, we surgically implanted a 100-electrode array with 1.5 mm shaft length (Blackrock Microsystems, Salt Lake City) in the primary motor cortex (M1) of each subject. We placed the subjects under isoflurane anesthesia and opened a small craniotomy above the motor cortex. We localized primary motor cortex using both visual landmarks and intracortical microstimulation to identify the arm region in Subjects M and C, and the hand region in Subject J. The arrays were then inserted pneumatically. During the behavioral experiments, we acquired neural data using a Blackrock Microsystems Cerebus system. The cortical signals were amplified and band-pass filtered (250 to 5000 Hz). To record the spiking activity of single neural units, we identified threshold crossings of six times the root-mean square (RMS) noise on each of the 96 recording channels and recorded spike times and brief waveform snippets surrounding the threshold crossings. For the first experiment, we recorded kinematic data from the robot handle at 1000 Hz using encoders in the manipulandum. For the second experiment, we recorded force data from the isometric box at 2000 Hz using the force-torque sensor. After each session, we sorted the neural waveform data using Offline Sorter (Plexon, Inc, Dallas, TX) to identify single neurons and discarded all waveforms believed to be multi-unit activity.

### Data preparation

After sorting the spike data, we estimated the firing rate of a neuron by binning the spike trains into non-overlapping windows of equal size. For the reaching task, we observed sparse firing rates and thus, we bin the data into 200 ms time bins and use data from the time that the ‘Go’ cue is given, to the end of the trial. For the isometric task, we used 50 ms time bins and restricted our analysis to a small time window immediately following the ‘Go’ cue (50 – 200 ms after the cue). The main neural dataset used in the paper (Subject M) contained *d* = 164 neurons and *T* = 1027 time points (across all eight reach directions). The dataset in Fig. 4 from Subject J, contained *d* = 67 neurons and *T* = 550 time points. To test the generalization of DAD to training sets from different subjects, we also used kinematics samples from Subject C, resulting in a dataset of size *T* = 803.

### Creating asymmetric tasks

For all of our experiments, we subsampled both datasets by removing directions of movement or force-production to produce datasets that were non-isotropic (asymmetric and non-reflective) and thus more representative of everyday tasks. To do this, we examined embeddings of the data to find directions of movement that are well represented by the neural data. Once we found these directions, we then sub-selected the entire movement and neural data to use only the trials in the selected directions. This approach is necessary to produce distributions that can be aligned with DAD.

### Basics of movement decoding

Before diving into DAD, we first need to define the variables for our decoding problem. Let **y**_*i*_ ∈ ℝ^*d*^ denote the firing rate of *d* neurons at the *i*^th^ point in time (sample). The time-varying neural activity of a population of *d* neurons can be represented as a matrix **Y** of size *d* × *T* by stacking the neural activity vectors {**y**_1_, **y**_2_, …, **y**_*T*_} into the columns of **Y**. The aim of movement decoding is to make use of the measured firing rates of a population to estimate the intended velocity vector 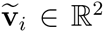 at the *i*^th^ point in time, where each entry of 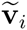 corresponds to the Cartesian velocities of movement in the *xy*-plane. Ultimately, our objective will be to map the neural activities to the space of decoded movements.

A large body of work has demonstrated that the relationship between the instantaneous velocity vector 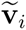 and the neural activity signal **y**_*i*_ in the motor cortex area M1 is approximately linear^29–31^. More recently, studies have shown that neurons are also tuned to to the magnitude of the 2D velocities^13^. Thus the kinematics and their magnitude, at the *i*^th^ point in time, can be decoded as **v**_*i*_ = **Hy**_*i*_, where **H** is a 3 × *d* matrix and **v**_*i*_ ∈ ℝ^3^ includes the magnitude of the kinematics as its third dimension.

In general, the right matrix for decoding (**H**) is unknown and must be estimated from neural and kinematics recordings. Here we assume that this linear model is time-invariant, and thus we can also write this model in matrix form as

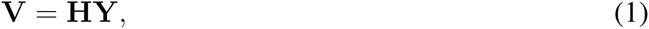

where **V** is a 3 × *T* matrix that contains the kinematics associated with the observed neural activities. When we have access to simultaneous recordings of kinematics **V** and the neural activity **Y**, this information can be used to estimate the matrix **H** (Eqn. 1). One way of solving this problem is to find a regularized least-squares estimate which minimizes the following loss function

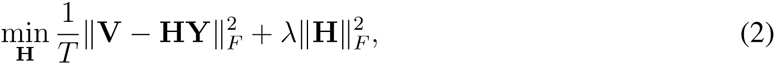

where 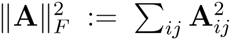 is the Frobenius norm and *λ* ≥ 0 is a user specified regularization parameter. This loss function defines the interplay between an error term and a term favoring simpler solutions.

### Distribution alignment decoding (DAD) approach

In contrast to the standard paradigm described above, we now consider the setting where we acquire *N* samples of the kinematics **V** ∈ ℝ^3×*N*^ separately from neural activity. Take for instance the case where we have recorded the kinematics from multiple users doing the same task and then, at a later time, we collect neural activity **Y** ∈ ℝ^*d*×*T*^ from a new user performing the same task. Since the kinematics **V** and neural activity **Y** are recorded at different times and are potentially of unequal dimension, determining the linear model **H** cannot be done by solving Eqn. 1. Even if the datasets contain the same number of samples (i.e. *N* = *T*), finding correspondence between the columns of **V** and the columns of **Y** is a NP hard problem^32^.

Rather than tackling the problem of finding correspondence between the two datasets, we can leverage the fact that the underlying distributions of samples in both spaces have similar structures to find a linear mapping that aligns the two. A natural framing of this problem is to find the best linear embedding of the neural data which minimizes the KL-divergence between the predicted distribution of kinematics (*q*) and the prior distribution of recorded kinematics (*p*). More formally, assume that the set of samples {**v**_1_, **v**_2_, …, **v**_*N*_} are drawn from a distribution *p* and that the pre-dicted kinematics vectors 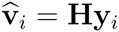 are drawn from a distribution *q*. To estimate **H**, our goal is to find a solution to the following optimization problem:

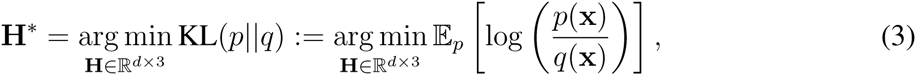

where the random variable **x** *∈* ℝ^3^ is drawn from the distribution *p*. By minimizing the KL-divergence, our approach essentially finds an affine transformation which maximizes the similarity between the distribution of transformed neural activity and prior distribution of kinematics.

In general, solving the problem in Eqn. 3 is intractable, due to the fact that the KL-divergence can be a non-convex and non-differentiable function of **H**. However, we can exploit the fact that, without substantial loss of information, **Y** can be projected into a lower-dimensional space where solving this problem is possible. We denote the resulting projected neural activities and the corresponding (low-d) decoder as **Y**_*ℓ*_ and **H**_*ℓ*_, respectively. When the mapping between **Y** and **V** is linear, the problem of distribution alignment can be reduced to finding the best linear transformation **H**_*ℓ*_ such that the KL-divergence between the distribution of observed kinematics *p* and predicted kinematics *q* is minimized. In a lower dimensional space, the alignment between the distributions of neural data and kinematics is feasible. To solve our low-dimensional alignment problem, DAD combines the benefits of fast closest point matching strategies^33^ with the use of the KL-divergence as a metric for alignment.

### Computing the KL-divergence

To measure the KL-divergence between the distributions *p* (target) and *q* (source) in Eqn. 3, we adopt a non-parametric approach that allows us to obtain an accurate estimate of these distributions without making any restrictive assumption on the form of either distribution. In particular, we rely on the popular *k*-nearest neighbors (KNN) density estimation algorithm^34,35^, which estimates a distribution using only distances between the samples and their *k*^th^ nearest neighbor. To make this concrete, we define the distance between a vector x and a matrix **A** as *ρ*_*k*_(**x**, **A**) = ||**x** – **a**_*k*_||_2_, where **a**_*k*_ is the *k*^th^ nearest neighbor to x contained in the columns of **A**. The value of the empirical distribution *p* at **v**_*i*_ is then estimated using the following consistent estimator^34^:

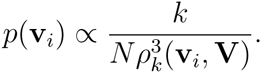

We partition the 3D space and then compute this quantity for every grid point. After computing the 3D density, we normalize to obtain a proper probability distribution over the space of grid points. Note that the intuition behind this approach is that in regions where we have higher density of samples, the k-nearest distance *ρ*_*k*_(**v**_*i*_, **V**) will be small and thus, the probability of generating a sample at this location is large. In practice, we set 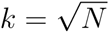 which guarantees that the estimates of *p* will asymptotically converge to the exact point estimates of the distribution since *ρ*_*k*_ converges to 0 as *N* → +∞ ^34^.

To estimate the distribution of the predicted kinematics 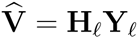 from the projected neural activities **Y**_*ℓ*_, we again use the same nearest neighbor approach. Assuming that we have an estimate for **H**_*ℓ*_ and **Y**_*ℓ*_, the empirical distribution of the predicted dynamics is given by 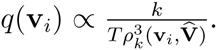. As with *p*, we compute this quantity for all *D* grid points and then normalize to produce a probability distribution of these points.

Once we compute the empirical distributions *p* and *q* over a fixed 3D grid {s_1_, …, s_*D*_}, we can use the following formula to estimate the KL-divergence:

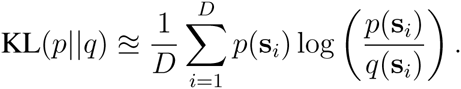

Upon estimating the probability distributions *p* and *q*, our aim is to find the best **H**_*ℓ*_ that minimizes their divergence. Solving this problem is intractable for large dimensions since the computational cost of solving Eqn. 3 scales exponentially with the dimension of the problem. Luckily, since the dimension of the task is small, it is possible to efficiently solve this optimization problem.

### Alignment algorithm

After reducing the dimensionality of the neural data, our alignment method consists of two main steps: 3D alignment (using a cone prior to regularize our problem) and then a fine-scale alignment in 2D. Both steps rely on the KL-divergence to measure the goodness of fit.

To align the datasets in 3D, we start by applying a 3D rotation to the source distribution (projected neural data) to initialize our search. After applying a 3D rotation, we apply an iterative closest point (ICP) algorithm^33^ to quickly match the two sets of points. This ICP step is important for finding the translation of the point clouds needed to obtain alignment. After aligning the datasets with ICP, we measure the KL-divergence between the target distribution and the alignment (when initializing with a 3D rotation). We repeat these steps over the space of 3D rotations (coarse grid search is sufficient) and measure the KL-divergence between each possible rotation and alignment. The alignment with minimal KL-divergence is selected as the output of our 3D alignment procedure.

After finding a 3D alignment, we often need to refine the estimate further. Thus we select the first two coordinates from our 3D embedding, which if the 3D alignment is done correctly, maps the data into the correct plane where the movement directions are visible. We then apply a similar alignment procedure in 2D, where we perform an exhaustive search over the rotation angle (2D) from 0-360 degrees and over two scaling parameters to define the skew of the data. We finally return the 2D embedding that minimizes the KL divergence between the target and source data.

### Dimensionality reduction

To quickly assess the low-dimensional structure of the data, we designed a module to pre-processes the neural data and apply a variety of dimensionality reduction techniques to the data. Using a Matlab toolbox for dimensionality reduction^15^, we applied PCA, factor analysis (FA), Isomap^36^, tSNE, SNE, Probabilistic PCA, and others. In addition, we also applied a Matlab toolbox for neural data analysis which implements generalized PCA^16^, using an exponential link function. We applied these techniques to our data and visually inspected the resulting embeddings to ensure that the task structure is indeed visible in the data. Note that this step can be done in a completely unsupervised way. After triaging the data to find embeddings with clear low-dimensional structure, we then applied DAD to align the neural data onto a prior movement distribution for the task. In our evaluations, we used factor analysis (FA) for all of the real motor datasets and used PCA for our synthetic experiments, as we found the more accurate and consistent embeddings with these methods.

### Incorporating prior knowledge into our alignment procedure

Due to the non-convexity of our KL-divergence minimization problem (Fig. 1b), it is important to use our prior knowledge about the shape of the target distribution in our optimization procedure^37^. We thus leverage the fact that the support of the distribution (regions with non-zero probability) lie near a cone in 3D. We confirm this shape both in the 3D projections of neural activity and also in the target kinematics when we take into account the magnitude of 2D kinematics as a third covariate in our linear model (Fig. 1d). Thus we can use this fact to improve the accuracy and efficiency of our alignment procedure by regularizing our KL-divergence problem and minimizing the number of possible local minima in our search. In practice, this means we leverage the bounded support of the distribution to appropriately rescale and then find a translation and scaling for distribution alignment.

### Using noise to improve shape matching

Our approach for shape matching via KL-divergence minimization seems to hold up well against synthetic data. However, when dealing with real neural datasets, we are sometimes faced with that problem that our target (clean) and source (messy) distributions do not look similar enough. Thus, when the size of the datasets are not matched, we create a blurred and oversampled version of our (clean) target kinematics data by replicating each data point and adding a small amount of Gaussian noise to each of these replicas. Our blurring procedure can be used to produce a better match between the clean target and messy source distributions.

### Synthetic model

When simulating neural activity, the firing rate of the *n*^th^ neuron at the *t*^th^ time point is given by

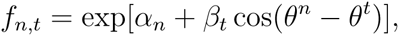

where *θ*^*n*^ is the preferred direction of the *n*^th^ neuron, *θ*^*t*^ = tan^*−*1^(*v*_*x,t*_/*v*_*y,t*_) is the direction of the movement at the *t*^th^ time point, *v*_*x,t*_ is the velocity in the x-direction at time t, *v*_*y,t*_ is the velocity in the y-direction at time t, and *α*_*n*_ and *β*_*n*_ are scalars which shift and modulate the firing rate, respectively. To generate spikes for a population of size *N*, we generate Poisson random variables according to the firing rates {*f*_1_, …, *f*_*N*_}. In our simulations (Fig. 4), we set the baseline *α*_*n*_ = 2, for all neurons (*∀n*) and set the modulation term 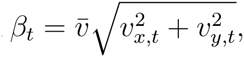 where 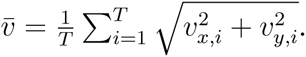

### Code availability and reproducibility

All of the code and scripts to generate figures are located at http://github.com/KordingLab/DAD. To generate the figures in this paper, we used neural data from Subject M and J, and motor variables from Subject’s M, C, and J. When applying DAD, there are a few parameters that we vary when processing the data and use later in alignment. The parameters are passed into the opts structure into the main wrapper function ‘runDAD.m’.

- Size of grid search: This parameter controls the number of 3D rotations to test as initial points in the 3D alignment step. We observe that a relatively coarse search is fine, we set this parameter to 3 (to generate 27 different angles in 3D) in all of the analysis in this paper.
- Noise variance: This parameter controls the variance of any noise added to the training distribution. We need to set this when the neural embeddings are very noisy and the matching procedure is not effective enough. In our experiments on Subject M, we do not add noise to the training data. For Subject J’s data, we often need to add some amount of noise—the results in Fig. 4 were generated by setting the variance to 0.1.
- Dimensionality reduction: Our code allows the user to specify the different dimensionality reduction methods they wish to test. To produce the results in the paper, we used factor analysis (FA) for all of our experiments on real data and PCA for our synthetic experiments.

